# Comparative clinical transcriptome of *pir* genes in severe *Plasmodium vivax* malaria

**DOI:** 10.1101/2025.07.10.664071

**Authors:** Pon Arunachalam Boopathi, Saurabh Singh, Priyanka Roy, Sampreeti Tahbildar, Sanjay Kumar Kochar, Dhanpat Kumar Kochar, Ashis Das

## Abstract

*vir* genes, a multigene family in *Plasmodium vivax* that are a part of a larger superfamily of genes called the *pir* (*Plasmodium* interspersed repeat) genes, have been reported earlier to be potentially involved in cyto-adherence and evasion of splenic clearance. *Plasmodium vivax*, historically characterised as a “benign” malaria parasite, has been associated with clinical outcomes including hepatic dysfunction, renal failure, and cerebral malaria in India and several global regions. It constitutes an economic burden and presents a public health challenge alongside other *Plasmodium* species. Here, we present a part of global transcriptomic studies using custom-designed microarrays that compare the transcriptome of the parasite responsible for severe *Plasmodium vivax* manifestations, specifically hepatic dysfunction and cerebral malaria from India, with an emphasis on the *pir* genes, some of which are reported to play a role in cyto-adherence. The RNA of the parasite isolated from 23 patients (uncomplicated group = 6, hepatic dysfunction group = 12, and cerebral malaria group = 5) was subjected to microarray hybridisation, and the data obtained showed a wide range of *pir* subfamilies to have been differentially expressed. We report the upregulation of 24 *pir* genes in the cerebral malaria group (n = 5) and 28 *pir* genes in the hepatic dysfunction group (n = 12), which belong to different subfamilies in at least 50% of the severe malaria patients’ group. Out of the upregulated *pir* genes in the cerebral malaria group, members of *vir* subfamily E (n=8 genes) and the *pvpir* subfamily H (n=6 genes) are expressed in a higher proportion compared to the hepatic dysfunction group, where members of *vir* subfamily E (n=9) and C (n=6) are expressed in a major proportion.

**Author Summary:** *Plasmodium vivax*, considered to cause benign infections, has been reported over the past couple of decades as manifesting with severe symptoms like cerebral malaria, hepatic dysfunction, and others. This paper reports the analysis of a selected *pir* gene repertoire from microarray hybridisation performed with custom-designed 15K microarrays. Differences between *pir* genes, which are specifically upregulated in either hepatic dysfunction or cerebral malaria, the two manifestations considered in addition to uncomplicated *vivax* malaria samples, are highlighted. The latter provided the control values for comparison with the disease groups. The data presented in this study show differentially upregulated *pir* genes, some of which are also common to both manifestations. Literature has reported one of the VIR proteins, encoded by *vir*14 C gene, to be involved in adhesion to ICAM-1. The future scope of this work includes validation with a larger sample size and characterising the products of the *pir* gene, as depicted with reference to their role in cytoadherence or other mechanisms that could lead to severe disease. It would be possible to devise strategies based on this data that could lead to the use of some of these molecules as potential biomarkers or for therapeutic intervention.

## Introduction

Malaria, an important vector-borne disease, continues to pose a major global public health challenge (1). Five parasitic protozoa species from the genus *Plasmodium*: *Plasmodium falciparum, Plasmodium vivax, Plasmodium malariae, Plasmodium ovale,* and *Plasmodium knowlesi* are responsible for this disease in humans (2)*. P. vivax* is the second most common cause of malaria globally, after *P. falciparum* (3). More than half of all cases (approx. 2.7 million) in the WHO South-East Asia region are due to *P. vivax* (1). In the past*, P. vivax* malaria was considered benign, particularly in comparison to *P. falciparum* malaria. Over the past twenty years, there has been a significant increase in the reports of severe disease manifestations caused by *P. vivax* mono-infections (4–13). Severe disease manifestations by this parasite include cerebral malaria (CM), hepatic dysfunction (HDYS), renal complications and others, as per the World Health Organisation criteria for severity (14).

In *P. vivax*, a multigene family known as the *vir* (15) has been linked to the parasite’s capacity for immune system escape and cytoadherence (16). These *vir* genes are a part of a larger superfamily of genes called the *pir* (*Plasmodium* interspersed repeat) genes (17). Initially, 346 *vir* genes were identified in the genome of *P. vivax*; these genes were primarily located in subtelomeric regions and were categorised into 12 subfamilies (18). Later, with the help of computational approaches, this multigene family was redefined into *vir* genes (B, C, E, G, I, J, K, and L subfamily); the newly defined *pvpir* genes (A, D and H subfamily); and the unclustered (not clustered) and unclassified genes (19). Over 1000 *pir* genes have been found in *P. vivax* genomes thus far (20). Expression studies of these *pir* genes revealed that individual isolates from patients expressed a large number of pir genes from various subfamilies (19,21,22). This vast diversity of PIR antigens has been suggested as allowing the parasite to evade the host immune system (21) and some of these have also been hypothesised to aid the parasite’s survival from splenic clearance (22). Studies on a few of the VIR proteins showed that they may have distinct subcellular locations and, thus, distinct roles. PVX_108770 of the *vir* subfamily C (23) and Pv11101_18 of the *vir* subfamily E (24) may be involved in mediating the parasite’s cyto-adhesion phenotype.

Microarrays and RNA Sequencing are two examples of high-throughput technologies frequently used to study the genetic responses that underlie host-parasite interactions. Microarrays have been utilised in the past to investigate various biological processes in *Plasmodium* spp. (25,26) and have now become a universal tool for studying the expression of thousands of genes (27,28). In our earlier work, we conducted a series of custom-designed 15K microarray experiments (29) on *Plasmodium vivax* field isolates obtained from patients presenting with either severe or uncomplicated malaria. In this study by *Boopathi et al* (29), a total of 267 *pir* genes were detected, while 259 *pir* genes were identified in the present analysis; notably, all 259 *pir* genes detected here were also reported in our earlier work. However, the primary objective of the previous study was the development, validation, and application of the *P. vivax* gene microarray, and differential expression analysis was performed using group-level mean expression values comparing complicated malaria (n = 9) with uncomplicated malaria (n = 5).

In contrast, the current study employs an individual-level transcriptomic analysis, with separate comparisons conducted for each clinical isolate across two independent disease categories: (i) cerebral malaria versus uncomplicated malaria and (ii) hepatic dysfunction versus uncomplicated malaria. This analytical strategy allows a more refined characterization of *pir* gene expression patterns and captures inter-individual variability that may be obscured by group-averaged approaches. Building on these findings, and motivated by the emerging evidence highlighting the extensive *pir* genes as a potential contributor to *P. vivax* pathogenesis, the present study represents the first comprehensive analysis of *pir* superfamily gene expression patterns in *P. vivax* isolates associated with severe clinical manifestations, specifically cerebral malaria (CM) and hepatic dysfunction (HDYS). Notably, the availability of patient blood samples from *P. vivax* cases presenting with CM or HDYS is inherently limited, owing to the relatively low incidence of these severe manifestations compared to uncomplicated *P. vivax* infections, underscoring the uniqueness and clinical relevance of this dataset.

## Materials and methods

### Patient details

We obtained blood samples from 23 patients with malaria caused by *P. vivax*. All samples were collected before drug treatment. The categorisation of severe malaria into cerebral malaria and hepatic dysfunction cases was according to WHO guidelines (WHO, 2015, 2010) (14). The patients were diagnosed as either severe (n=17; PVC-1, PVC-2, PVC-8, PVC-20, PVC-21, PVC-22, PVC-25, PVC-27, PVC-28, PVC-30, PVC-31, PVC-32, PVC-11, PVC-15, PVC-16, PVC-17 & PVC-18) or uncomplicated malaria (n=6; PVU-2, PVU-3, PVU-6, PVU-9, PVU-10 & PVU-16). Under the severe category (n=17), 12 were jaundice/hepatic dysfunction cases, and 5 were cerebral malaria cases. Two patients from the uncomplicated malaria group (PVU-9 and PVU-10) and eight patients from the HDYS group (PVC-1, PVC-2, PVC-20, PVC-21, PVC-27, PVC-28, PVC-30, and PVC-31) are the same as described in *Boopathi et al* (29). However, in the cited manuscript, the analysis was performed differently from the way it is reported in this paper. Clinical profile of all the patients included in this study are presented in the Table 1. All cases of hepatic dysfunction had a serum bilirubin level greater than 3 mg/dL, and all cerebral malaria cases had a Glasgow Coma Scale score ranging from 7 to 9, as per WHO criteria. All patients with severe malaria underwent appropriate blood tests to rule out viral hepatitis, dengue, typhoid, leptospirosis, infectious mononucleosis, and HIV. All the tests were negative, indicating that the reported symptoms were solely due to *P. vivax* infection.

**Table 1.**
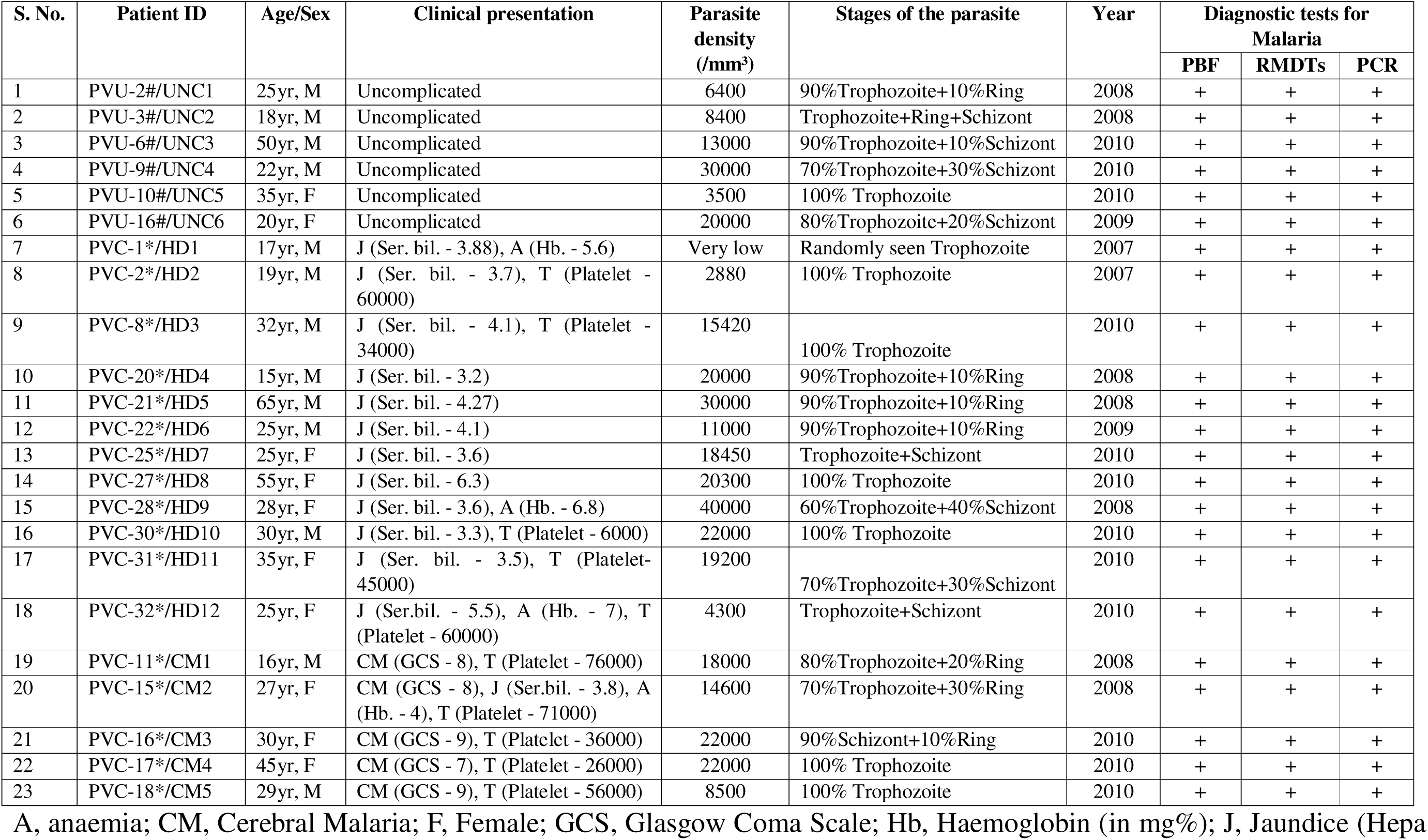
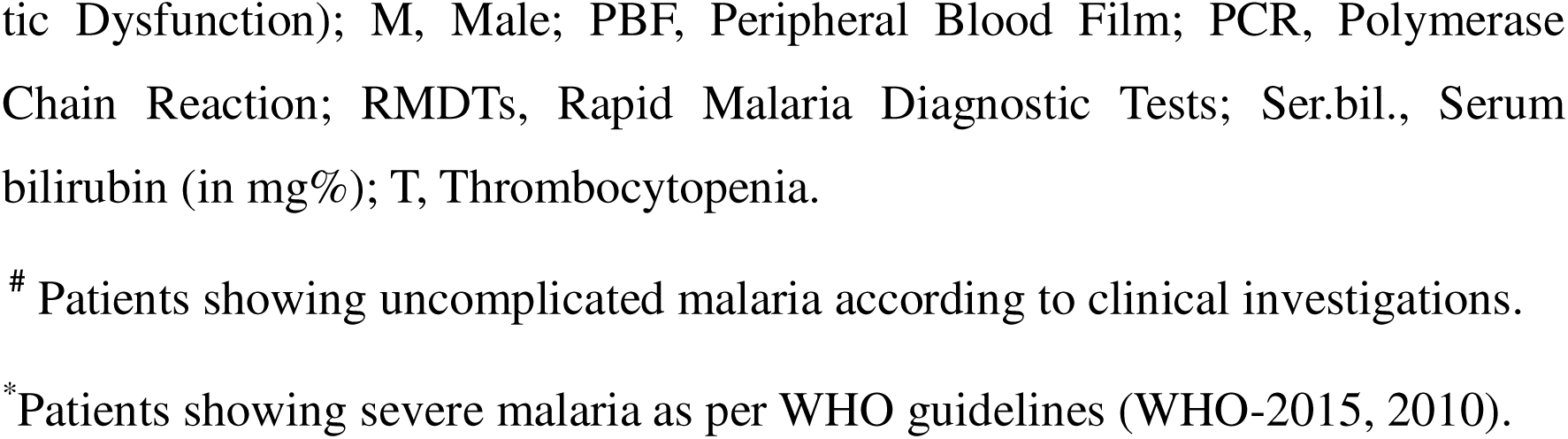
Clinical characteristics of patients infected with *P. vivax*.

| S. No. | Patient ID | Age/Sex | Clinical presentation | Parasite density (/mm<sup>3</sup>) | Stages of the parasite | Year | Diagnostic tests for Malaria |  |  |
| --- | --- | --- | --- | --- | --- | --- | --- | --- | --- |
|  |  |  |  |  |  |  | PBF | RMDTs | PCR |
| 1 | PVU-2#/UNC1 | 25yr, M | Uncomplicated | 6400 | 90% Trophozoite+10% Ring | 2008 | + | + | + |
| 2 | PVU-3#/UNC2 | 18yr, M | Uncomplicated | 8400 | Trophozoite+Ring+Schizont | 2008 | + | + | + |
| 3 | PVU-6#/UNC3 | 50yr, M | Uncomplicated | 13000 | 90% Trophozoite+10% Schizont | 2010 | + | + | + |
| 4 | PVU-9#/UNC4 | 22yr, M | Uncomplicated | 30000 | 70% Trophozoite+30% Schizont | 2010 | + | + | + |
| 5 | PVU-10#/UNC5 | 35yr, F | Uncomplicated | 3500 | 100% Trophozoite | 2010 | + | + | + |
| 6 | PVU-16#/UNC6 | 20yr, F | Uncomplicated | 20000 | 80% Trophozoite+20% Schizont | 2009 | + | + | + |
| 7 | PVC-1*/HD1 | 17yr, M | J (Ser. bil. - 3.88), A (Hb. - 5.6) | Very low | Randomly seen Trophozoite | 2007 | + | + | + |
| 8 | PVC-2*/HD2 | 19yr, M | J (Ser. bil. - 3.7), T (Platelet - 60000) | 2880 | 100% Trophozoite | 2007 | + | + | + |
| 9 | PVC-8*/HD3 | 32yr, M | J (Ser. bil. - 4.1), T (Platelet - 34000) | 15420 | 100% Trophozoite | 2010 | + | + | + |
| 10 | PVC-20*/HD4 | 15yr, M | J (Ser. bil. - 3.2) | 20000 | 90% Trophozoite+10% Ring | 2008 | + | + | + |
| 11 | PVC-21*/HD5 | 65yr, M | J (Ser. bil. - 4.27) | 30000 | 90% Trophozoite+10% Ring | 2008 | + | + | + |
| 12 | PVC-22*/HD6 | 25yr, M | J (Ser. bil. - 4.1) | 11000 | 90% Trophozoite+10% Ring | 2009 | + | + | + |
| 13 | PVC-25*/HD7 | 25yr, F | J (Ser. bil. - 3.6) | 18450 | Trophozoite+Schizont | 2010 | + | + | + |
| 14 | PVC-27*/HD8 | 55yr, F | J (Ser. bil. - 6.3) | 20300 | 100% Trophozoite | 2010 | + | + | + |
| 15 | PVC-28*/HD9 | 28yr, F | J (Ser. bil. - 3.6), A (Hb. - 6.8) | 40000 | 60% Trophozoite+40% Schizont | 2008 | + | + | + |
| 16 | PVC-30*/HD10 | 30yr, M | J (Ser. bil. - 3.3), T (Platelet - 6000) | 22000 | 100% Trophozoite | 2010 | + | + | + |
| 17 | PVC-31*/HD11 | 35yr, F | J (Ser. bil. - 3.5), T (Platelet - 45000) | 19200 | 70% Trophozoite+30% Schizont | 2010 | + | + | + |
| 18 | PVC-32*/HD12 | 25yr, F | J (Ser. bil. - 5.5), A (Hb. - 7), T (Platelet - 60000) | 4300 | Trophozoite+Schizont | 2010 | + | + | + |
| 19 | PVC-11*/CM1 | 16yr, M | CM (GCS - 8), T (Platelet - 76000) | 18000 | 80% Trophozoite+20% Ring | 2008 | + | + | + |
| 20 | PVC-15*/CM2 | 27yr, F | CM (GCS - 8), J (Ser. bil. - 3.8), A (Hb. - 4), T (Platelet - 71000) | 14600 | 70% Trophozoite+30% Ring | 2008 | + | + | + |
| 21 | PVC-16*/CM3 | 30yr, F | CM (GCS - 9), T (Platelet - 36000) | 22000 | 90% Schizont+10% Ring | 2010 | + | + | + |
| 22 | PVC-17*/CM4 | 45yr, F | CM (GCS - 7), T (Platelet - 26000) | 22000 | 100% Trophozoite | 2010 | + | + | + |
| 23 | PVC-18*/CM5 | 29yr, M | CM (GCS - 9), T (Platelet - 56000) | 8500 | 100% Trophozoite | 2010 | + | + | + |

A, anaemia; CM, Cerebral Malaria; F, Female; GCS, Glasgow Coma Scale; Hb, Haemoglobin (in mg%); J, Jaundice (Hepa
tic Dysfunction); M, Male; PBF, Peripheral Blood Film; PCR, Polymerase Chain Reaction; RMDTs, Rapid Malaria Diagnostic Tests; Ser.bil., Serum bilirubin (in mg%); T, Thrombocytopenia.

### Blood sample collection

Venous blood samples (∼5 mL) were collected from *P. vivax*-infected patients with their informed written consent at S.P. Medical College, Bikaner, India, between 2007 and 2010. Microscopy and rapid diagnostic tests (RDTs) for malaria were used for preliminary screening. Blood collected within 15 minutes was subjected to density gradient-based separation (Histopaque 1077, Sigma Aldrich, USA) to separate peripheral blood mononuclear cells (PBMCs) from infected and uninfected erythrocytes, in accordance with the manufacturer’s instructions. Both fractions were washed twice with phosphate-buffered saline (PBS) and then lysed with 4 volumes of Tri-reagent (Sigma Aldrich, USA). The samples were kept at -80 . Subsequently, samples were transported in a cold chain to BITS Pilani and processed. The isolated DNA of all the 23 malaria-infected patients were assessed by 18S rRNA-based multiplex PCR (30,31) to confirm *P. vivax* infection and rule out any chance of *P. falciparum* co-infection. The samples were collected under the ethical approval of the Institute Ethics Committee (IEC) of Sardar Patel Medical College, Bikaner, for sample collection (No. F. (Acad) SPMC/2003/2395) and permission to use these samples for related studies, as approved by Institute Ethics and Research Board (IERB) (No. F29(Acad)SPMC/2020/3151.

### 15K P. vivax microarray and experimentation

#### Sample preparation

Total RNA was extracted from severe (n=17) and uncomplicated (n=6) malaria patients’ blood samples, following the manufacturer’s protocol (Tri-Reagent, Sigma Aldrich, USA). The DNA was resuspended in 90 µl TE buffer and stored at -20°C. The RNA was resuspended in 90 µl of DEPC-treated deionised water and stored at -80°C. The quality of the RNA samples was assessed using the RNA 6000 Nano LabChip on the 2100 Bioanalyzer (Agilent, Palo Alto, CA) and formaldehyde-based denaturing agarose gel electrophoresis. The quality and quantity of the total RNA from the samples were evaluated separately using the NanoDrop ND-1000 UV-Vis Spectrophotometer (NanoDrop Technologies, Rockland, USA).

#### Microarray design and analysis

We developed a high-density, experimentally optimised 15K *P. vivax* oligonucleotide microarray (29). A total of 14342 validated probes representing 4180 and 2260 genes of *P. vivax* denoted in sense, and antisense orientation were chosen from the best performing probes, defined as one exhibiting a minimum two fold signal intensity above background, ensuring specificity and reliability in the microarray from a previously custom-designed 244K microarray (32). In our analysis, we considered the expression profile of *pir* genes in individual *P. vivax* samples with severe malaria. 267 *pir* genes, as classified by *Lopez et al* (19) were represented in this microarray. We applied a filtering criterion of transcript detection in at least 1 out of 11 samples in the cerebral malaria group (PVC = 5 and PVU = 6) and in at least 1 out of 18 samples in the hepatic dysfunction group (PVC = 12 and PVU = 6). A quantile normalisation with baseline to median of control samples (*P. vivax* uncomplicated malaria patients’ group; n=6), which resulted in differential expression profile of 260 *pir* genes in cerebral malaria samples and 259 *pir* genes in hepatic dysfunction cases (S1 Table). The log2 fold change value of +1 and above was considered as upregulated, and -1 and below was considered as downregulated. *Pir* genes upregulated in at least 50% across the severe malaria patients’ group (CM n=3/5 to 5/5; HDYS n=6/12 to 12/12) have been considered in this study for both manifestations. Raw data from this microarray experimentation were submitted to GEO and are available under the series ID GSE301003 (https://www.ncbi.nlm.nih.gov/geo/query/acc.cgi?acc=GSE301003). The data will be made publicly available if the manuscript is accepted.

### qPCR validation of microarray data

Quantitative real-time PCR (qPCR) was used to validate the microarray expression of a few selected genes. Primer3 (33) was utilised to create the primers required for this purpose. The details of the primers are provided in S1 Fig. DNA contamination was removed from the total RNA using the manufacturer’s protocol with the iScript gDNA Clear cDNA Synthesis Kit (BIO-RAD). The removal of DNA in RNA samples was verified by the absence of a DNA band in a 2% agarose gel after 40 cycles of PCR using Seryl-tRNA synthetase primers. 500ng of total RNA from uncomplicated (n=6) and severe (n=17) malaria patients’ samples was pooled at an equimolar concentration. Following the manufacturer’s protocol, first-strand cDNA synthesis was carried out using iScript gDNA Clear cDNA Synthesis Kit (BIO-RAD) in a total volume of 20µl. The Seryl-tRNA synthetase gene was taken as an endogenous control. The PCR conditions for all selected genes are provided in S1 Fig. Log2 fold change expression was calculated to determine the status of each gene.

## Results and Discussion

To understand the severity of disease caused by *P. vivax*, studying the expression profile of *pir* genes is important, as a few of the *pir* genes (from *vir* subfamily C and E) have been reported earlier to be possibly involved in cytoadherence and evasion of splenic clearance (22,34,35). *Pir* genes are poly-clonally expressed genes that have the potential to present proteins on the surface of iRBCs or localised elsewhere on the parasite’s surface or cytoplasm. These could have multiple roles in the biology of parasites. PIRs expressed on the surface of iRBCs, because of their inherent diversities, may be involved in immune evasion mechanisms in the host (22). On the other hand, the literature suggests an alternative role for VIR-based cytoadhesion, namely, evasion of splenic clearance during infection (35). It is possible that certain host or parasite factors may trigger the overexpression of a select group of PIR proteins, which then mediate cytoadhesion and other phenomena, leading to severe disease manifestations.

Here, in this study, with the help of a 15K custom-designed microarray, we analysed the difference in the expression pattern of *pir* superfamily members between patients with cerebral malaria (CM) and those with hepatic dysfunction (HDYS) manifestation from the Bikaner region of North-West India. Although, RNA sequencing can indeed provide a more comprehensive and sensitive approach for transcriptomic analyses. However, microarray-based platforms remain more cost-effective, particularly when working with clinical samples and larger sample sets. Additionally, RNA-seq typically requires high-quality RNA, to ensure reliable and unbiased results. RNA samples obtained *ex vivo* from enriched patient blood samples, such as those used in this study, may sometimes have RNA Integrity Number (RIN) slightly below the threshold (RIN 7). Under these circumstances, microarrays which rely on probe-based hybridisation are more tolerant of moderately reduced RNA integrity and therefore provide a practical and reliable alternative. These considerations influenced our decision to use microarray analysis in the present study.

The Sal-1 genome was used as the reference for our microarray design as it was the most comprehensive and widely available *P. vivax* genome at the time of study conceptualisation, and has been retained to ensure consistency and comparability with previously generated datasets. Although more recent Southeast Asian genomes (e.g., P01, W1) offer improved annotation, the relatedness of Rajasthan isolates to these strains remains unclear due to lack of comparative genomic data. Importantly, Sal-1 probes were empirically validated during 15K array development using a 244K platform, supporting their reliability. Future analyses incorporating multiple reference genomes may further improve resolution, particularly for the *pir* gene family.

### Analysis of the expression data of *pir* genes

On analysing upregulated *pir* genes, 24 *pir* genes in cerebral malaria patients’ group (n=5) and 28 *pir* genes in hepatic dysfunction patients’ group (n=12) belonging to different subfamily have been shown to be upregulated in at least 50% (CM n=3/5 to 5/5; HDYS n=6/12 to 12/12) of the severe malaria patients’ group (Figs 1 and 2) (S2 Table). Under cerebral malaria manifestation, members of the *vir* subfamily B, C, E, and G, as well as *pvpir* subfamily H and A, as well as unclassified and not clustered genes, have been upregulated in patient samples, ranging from 60-80% (n=3/5 to 4/5) (Fig 2) (S2 Table). Out of these 24 upregulated *pir* genes in cerebral malaria manifestations, the most prominent subfamilies were *vir* E (a third of the genes; n=8) and *pvpir* H (a quarter of the genes; n=6). Under hepatic dysfunction manifestation, members of the *vir* subfamily B, C, E, G, and I; *pvpir* subfamily H and A, as well as not clustered genes have been upregulated in the patient group, ranging from 50-91.7% (n=6/12 to 11/12) (Fig 2) (S2 Table). Out of the 28 upregulated *pir* genes in hepatic dysfunction manifestation, the most prominent subfamilies were *vir* E (n = 9) and C (n = 6).

**Fig 1.**
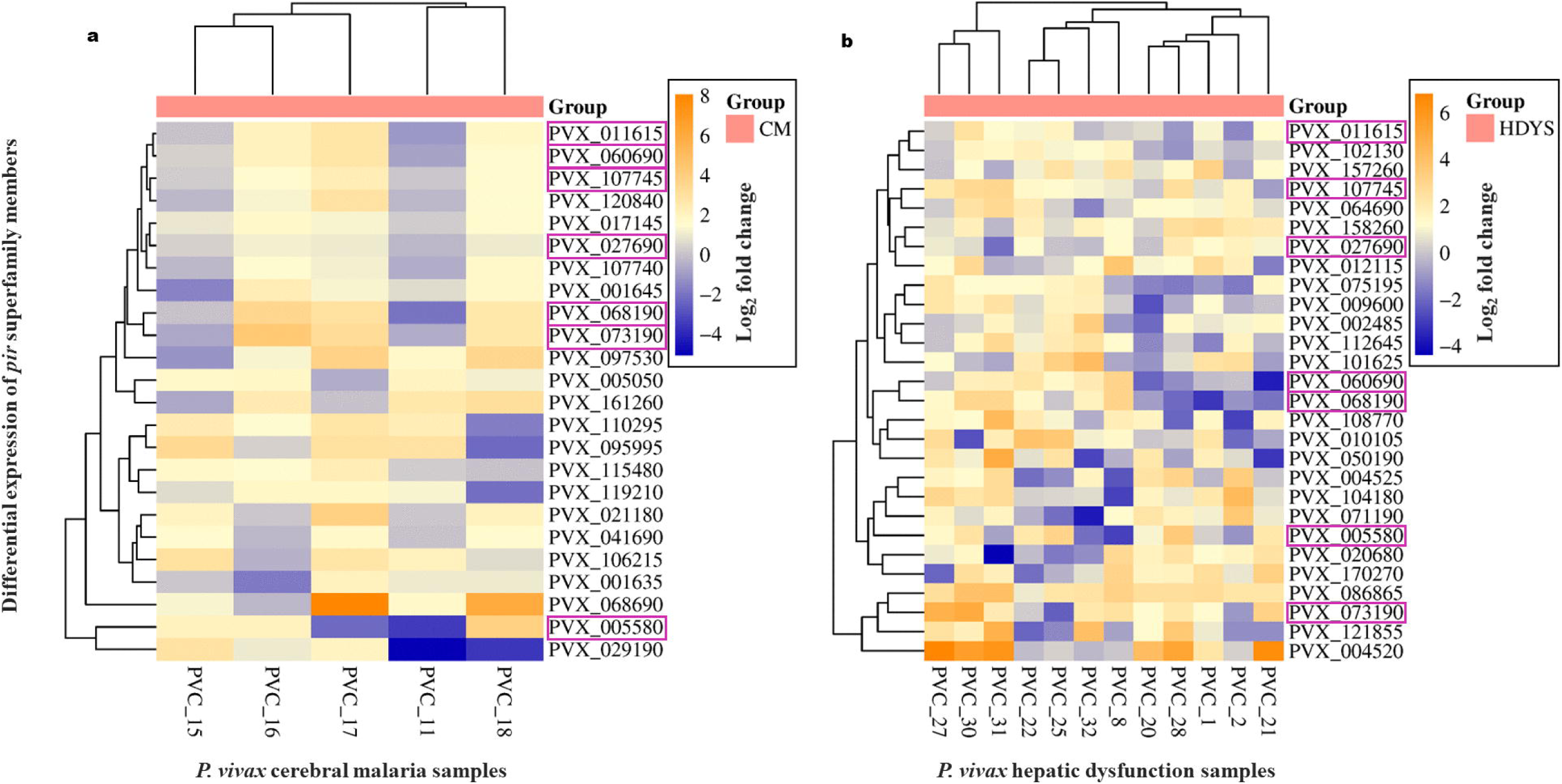
Clustered heat map expression profile of upregulated *pir* superfamily members in at least 50% (CM n=3/5 to 5/5; HDYS n=6/12 to 12/12) *P. vivax* severe malaria patients’ group (n=17) compared against the uncomplicated malaria patients’ group (n=6). (a) *P. vivax* cerebral malaria patients’ group (n=5). (b) *P. vivax* hepatic dysfunction patients’ group (n=12). The *pir* genes highlighted in magenta-colour outlined cells are common to both disease manifestations.

**Fig 2.**
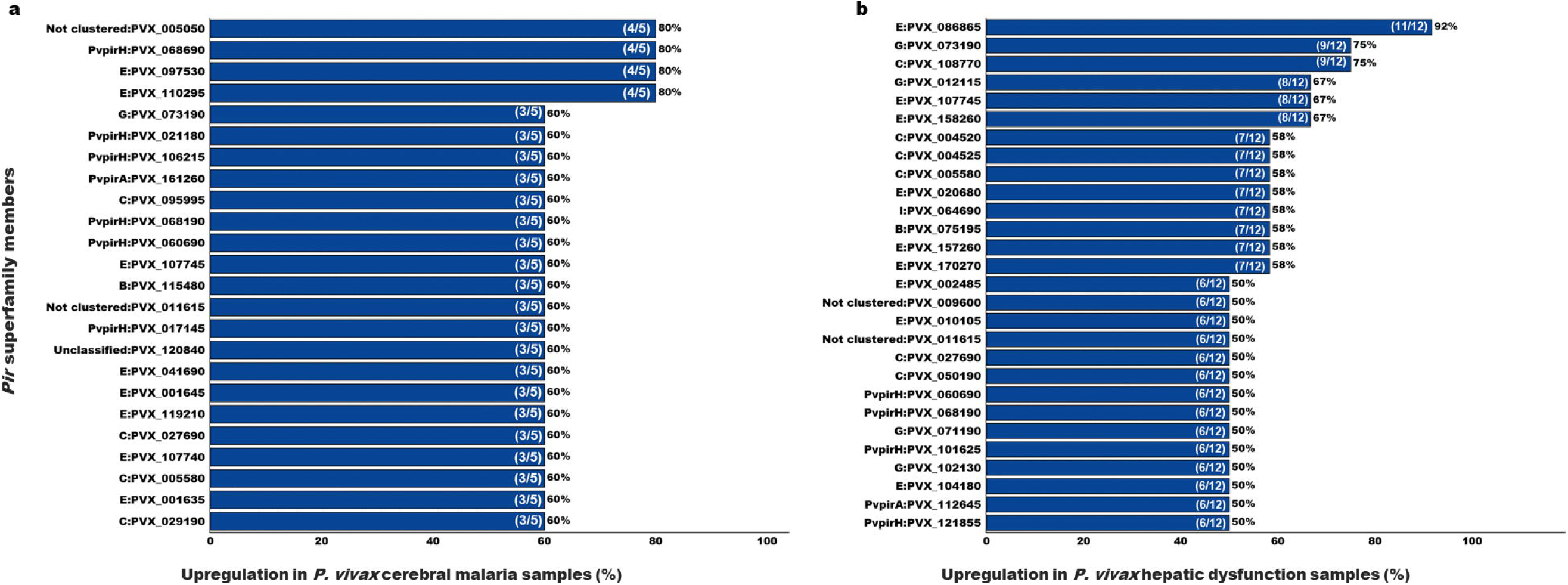
Upregulation of *pir* superfamily members in at least 50% (CM n=3/5 to 5/5; HDYS n=6/12 to 12/12) *P. vivax* severe malaria patient’s group (n=17) compared against uncomplicated malaria patients’ group (n=6). (a) *P. vivax* cerebral malaria patients’ group (n=5). (b) *P. vivax* hepatic dysfunction patients’ group (n=12).

The hierarchical clustering of *pir* genes, depicted in the heatmap (Fig 1), has aligned the genes according to their similar expression patterns within their respective patient groups. For example, in the cerebral malaria (CM) patient group, PVX_107745 (*vir* E) and PVX_120840 (Unclassified) are upregulated in PVC_16, PVC_17, and PVC_18, but are non-differentially expressed in PVC_11 and PVC_15. Similarly, in the hepatic dysfunction (HDYS) patient group, PVX_158260 (*vir* E) and PVX_027690 (*vir* C) are upregulated in PVC_1, PVC_2, PVC_8, PVC_21, PVC_22, and PVC_28, but are non-differentially expressed in PVC_20, PVC_25, and PVC_27.

On analysing downregulated *pir* genes, 28 genes in the cerebral malaria patients’ group (n=5) and 34 genes in the hepatic dysfunction patients’ group (n=12) belonging to different subfamilies have been shown to be downregulated in at least 50% (CM n=3/5 to 5/5; HDYS n=6/12 to 12/12) severe malaria patients’ group (S2 Table). Under cerebral malaria manifestation, members of the *vir* subfamily B, C, E, and G, as well as the *pvpir* subfamily H, as well as unclassified and not clustered genes, have been downregulated in the patient’s group, ranging from 60-100% (n=3/5 to 5/5) (S2 Table). Out of the 28 downregulated *pir* genes in cerebral malaria manifestation, the most prominent subfamilies were *vir* E (n = 14) and *pvpir* H (n = 4). Under hepatic dysfunction manifestation, members of the *vir* subfamily C, E, G, and K, as well as the *pvpir* subfamily D and H, as well as unclassified and not clustered pir genes, have been downregulated in patients’ groups, ranging from 50-91.7% (n=6/12 to 11/12) (S2 Table). Out of the 34 downregulated *pir* genes in hepatic dysfunction manifestation, the most prominent subfamilies were *vir* E (n = 22) and *pvpir* H (n = 3) (S2 Table).

Further, microarray results of four *pir* genes (PVX_086865 (*vir* E), PVX_097530 (*vir* E), PVX_017645 (*pvpir* D) and PVX_104180 (*vir* E) were also confirmed by real-time qPCR experimentation, shown in Fig 3. Real time qPCR experiment confirms upregulation of two *pir* genes (PVX_086865 in HDYS patients’ group (n=11/12) & PVX_097530 in CM patients’ group (n=4/5)) and downregulation of two *pir* genes (PVX_017645 in HDYS patient’s group (n=11/12) & PVX_104180 in CM patients’ group (n=5/5)) from microarray data (S1 and S2 Tables). Overall, real-time qPCR analysis validated the expression pattern of selected *pir* genes as seen in the microarray data.

**Fig 3.**
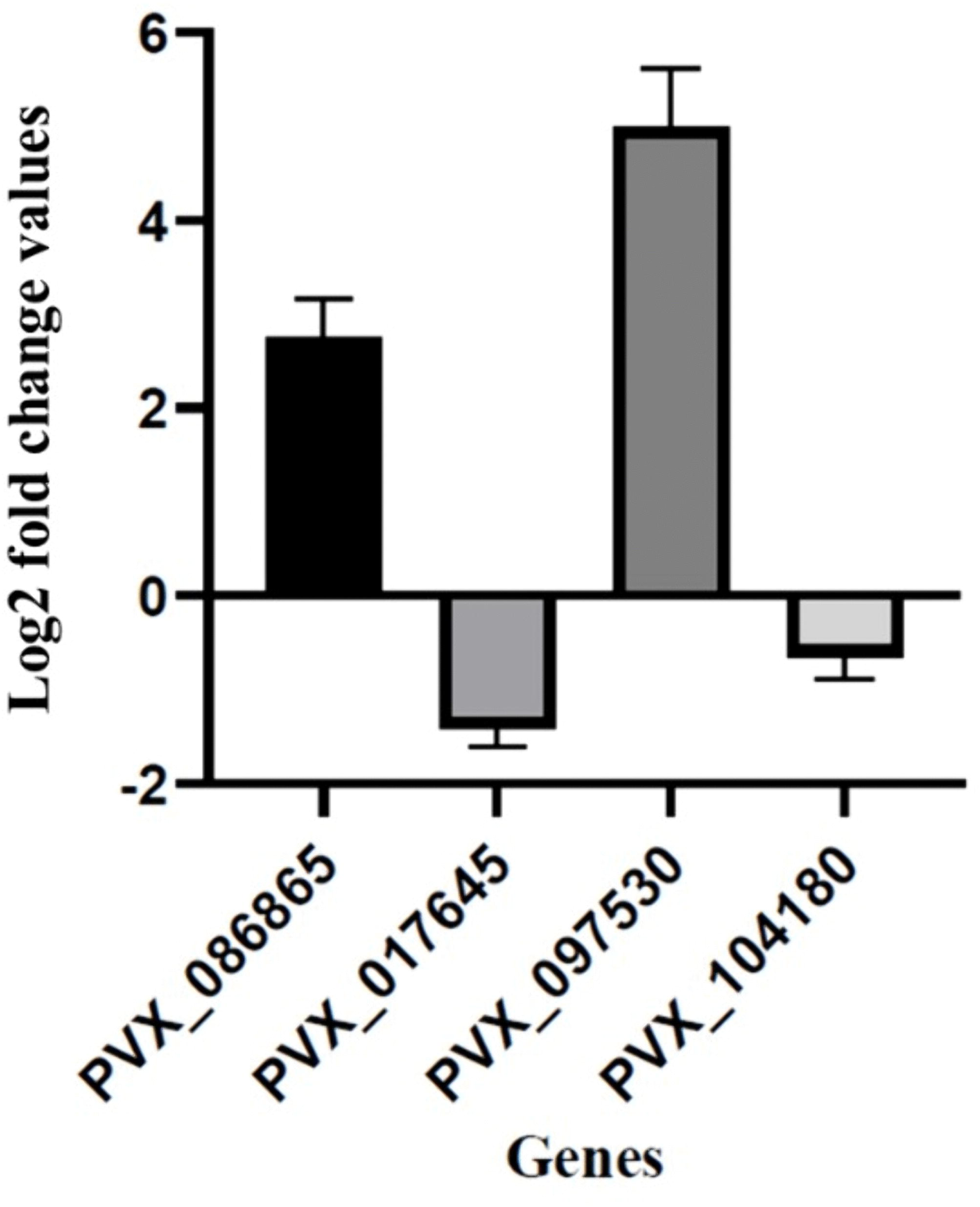
Validation of some *pir* genes by quantitative PCR (qPCR). Seryl tRNA synthetase was used as a reference gene. X-axis represents genes selected for validation; Y-axis represents Log2 fold change expression (mean ± corrected S.D.). PVX_086865, hypothetical protein (*vir* E); PVX_017645, variable surface protein Vir35 (*pvpir* D); PVX_097530, variable surface protein Vir22/12-related (*vir* E); PVX_104180, variable surface protein Vir12 (*vir* E).

### Comparison of the expressed *pir* genes between cerebral malaria and hepatic dysfunction cases

Out of the top at least 50% (CM n=3/5 to 5/5; HDYS n=6/12 to 12/12) upregulated *pir* genes in severe malaria patients’ group, 7 *pir* genes belonging to the *vir* subfamily C, G and E; *pvpir* subfamily H and not clustered are common among cerebral malaria and hepatic dysfunction cases (Fig 4a) (S2 Table). In cerebral malaria manifestation, 17 *pir* genes belonging to the *vir* subfamily B, C, and E; *pvpir* subfamily A and H; not clustered and unclassified are distinct (Fig 4a) (S2 Table). In hepatic dysfunction manifestation, 21 *pir* genes belonging to the *vir* subfamily B, C, E, G, and I; *pvpir* subfamily A and H; not clustered are distinct (Fig 4a) (S2 Table).

**Fig 4.**
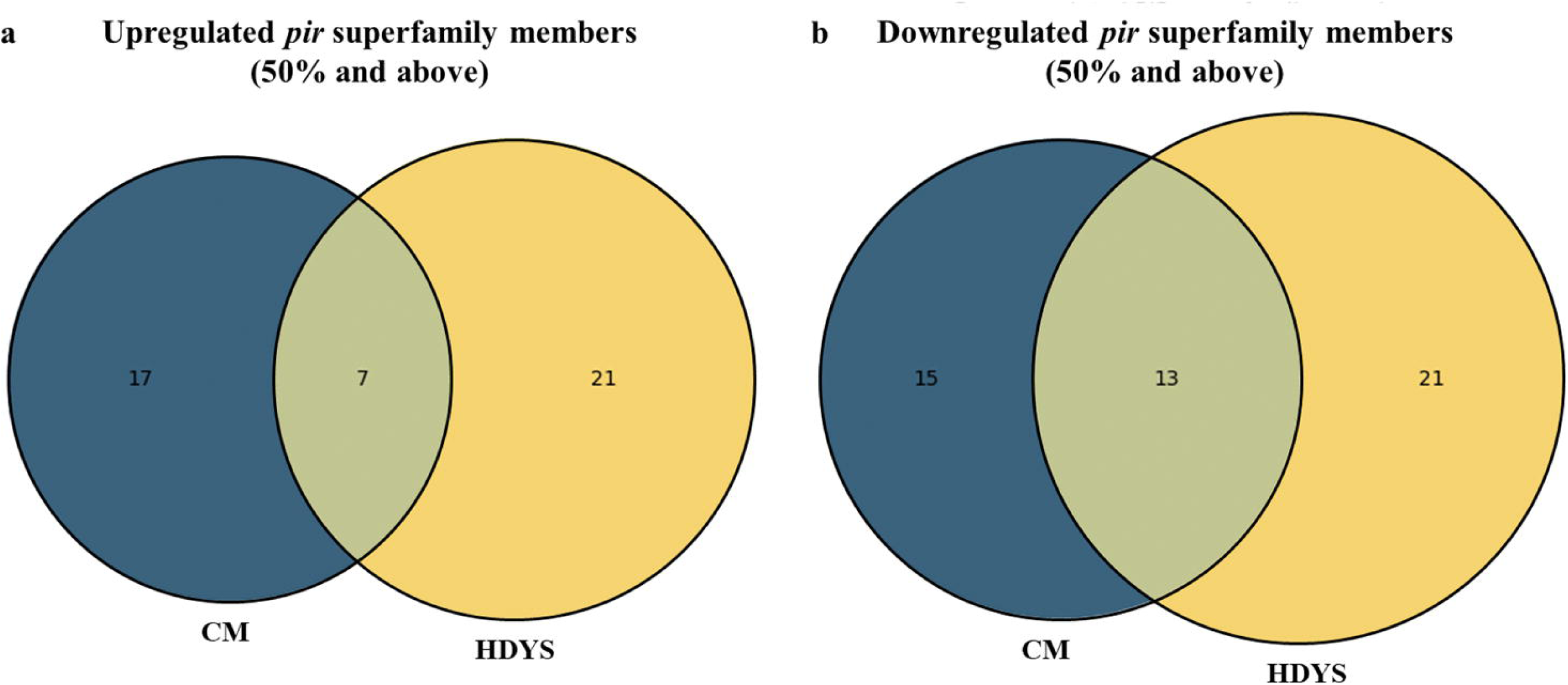
Distribution of *pir* superfamily members among cerebral malaria (n=5) and hepatic dysfunction (n=12) patients’ group. (a) At least 50% (CM n=3/5 to 5/5; HDYS n=6/12 to 12/12) upregulated *pir* genes. (b) At least 50% (CM n=3/5 to 5/5; HDYS n=6/12 to 12/12) downregulated *pir* genes.

Out of the top at least 50% (CM n=3/5 to 5/5; HDYS n=6/12 to 12/12) downregulated *pir* genes in severe malaria patients’ group, 13 *pir* genes belonging to the *vir* subfamily C, E and G; *pvpir* subfamily H; not clustered and unclassified are common among cerebral malaria and hepatic dysfunction cases (Fig 4b) (S2 Table). In cerebral malaria manifestation, 15 *pir* genes belonging to the *vir* subfamily B, C, E and G; *pvpir* subfamily H are distinct (Fig 4b) (S2 Table). In hepatic dysfunction manifestation, 21 *pir* genes belonging to the *vir* subfamily C, E, and K; *pvpir* subfamily D and H; not clustered are distinct (Fig 4b) (S2 Table).

A notable finding from our analysis is that the specific *pir* genes belonging to individual subfamilies, which have been upregulated in at least 50% (CM, n = 3/5 to 5/5; HDYS, n = 6/12 to 12/12) of the severe malaria patient group studied, are distinct between the disease manifestations (Table 2).

**Table 2.** Complication-specific *pir* gene upregulation.

| Upregulated *pir* gene description |  | Upregulation in *P. vivax* isolates showing **Hepatic Dysfunction** (n=12) 12PVC vs 6PVU |  | Upregulation in *P. vivax* isolates showing **Cerebral Malaria** (n=5) 5PVC vs 6PVU |  | Upregulated *pir* gene description |  | Upregulation in *P. vivax* isolates showing **Hepatic Dysfunction** (n=12) 12PVC vs 6PVU |  | Upregulation in *P. vivax* isolates showing **Cerebral Malaria** (n=5) 5PVC vs 6PVU |  |
| --- | --- | --- | --- | --- | --- | --- | --- | --- | --- | --- | --- |
| Gene Symbol | PIR subfamily | No. of samples | % | No. of samples | % | Gene Symbol | PIR subfamily | No. of samples | % | No. of samples | % |
| PVX_086865 | *vir* E | 11 | 92 | 2 | 40 | PVX_112645 | *pvpir* A | 6 | 50 | Nil | Nil |
| PVX_108770 | *vir* C | 9 | 75 | 1 | 20 | PVX_121855 | *pvpir* H | 6 | 50 | 1 | 20 |
| PVX_012115 | *vir* G | 8 | 67 | 2 | 40 | PVX_005050 | NC | 4 | 33 | 4 | 80 |
| PVX_158260 | *vir* E | 8 | 67 | 2 | 40 | PVX_068690 | *pvpir* H | 4 | 33 | 4 | 80 |
| PVX_004520 | *vir* C | 7 | 58 | 2 | 40 | PVX_097530 | *vir* E | 3 | 25 | 4 | 80 |
| PVX_004525 | *vir* C | 7 | 58 | 2 | 40 | PVX_110295 | *vir* E | 3 | 25 | 4 | 80 |
| PVX_020680 | *vir* E | 7 | 58 | Nil | Nil | PVX_001635 | *vir* E | 3 | 25 | 3 | 60 |
| PVX_064690 | *vir* I | 7 | 58 | 2 | 40 | PVX_001645 | *vir* E | 4 | 33 | 3 | 60 |
| PVX_075195 | *vir B* | 7 | 58 | 2 | 40 | PVX_017145 | *pvpir H* | 5 | 42 | 3 | 60 |
| PVX_157260 | *vir E* | 7 | 58 | 2 | 40 | PVX_021180 | *pvpir H* | 2 | 17 | 3 | 60 |
| PVX_170270 | *vir E* | 7 | 58 | 2 | 40 | PVX_029190 | *vir C* | 5 | 42 | 3 | 60 |
| PVX_002485 | *vir E* | 6 | 50 | Nil | Nil | PVX_041690 | *vir E* | 4 | 33 | 3 | 60 |
| PVX_009600 | NC | 6 | 50 | 1 | 20 | PVX_095995 | *vir C* | 5 | 42 | 3 | 60 |
| PVX_010105 | *vir E* | 6 | 50 | 1 | 20 | PVX_106215 | *pvpir H* | 4 | 33 | 3 | 60 |
| PVX_050190 | *vir C* | 6 | 50 | 1 | 20 | PVX_107740 | *vir E* | 3 | 25 | 3 | 60 |
| PVX_071190 | *vir G* | 6 | 50 | 1 | 20 | PVX_115480 | *vir B* | 1 | 8 | 3 | 60 |
| PVX_102130 | *vir G* | 6 | 50 | 1 | 20 | PVX_119210 | *vir E* | 3 | 25 | 3 | 60 |
| PVX_104180 | *vir E* | 6 | 50 | Nil | Nil | PVX_120840 | Unclassified | 5 | 42 | 3 | 60 |
| PVX_101625 | *pvpir H* | 6 | 50 | 1 | 20 | PVX_161260 | *pvpir A* | 4 | 33 | 3 | 60 |

■ At least 50% upregulation in the hepatic dysfunction group and less than 50% upregulation in the cerebral malaria group
■ At least 50% upregulation in the cerebral malaria group and less than 50% upregulation in the hepatic dysfunction group

Further looking into the expression profiles of distinct *pir* genes upregulated in one disease group and their downregulation status in the other disease group (S3 Table) only two *pir* genes (PVX_119210 (*vir* E) upregulated in CM group; PVX_104180 (*vir* E) upregulated in HDYS group) have been identified with opposite expression profiles (Upregulated in CM and downregulated in HDYS, or vice versa), as most of the distinct *pir* genes upregulated in one disease group are non-differentially expressed in the other disease group (S3 Table). Although these patterns do not allow for definitive conclusions regarding the strict opposite expression of *pir* genes between disease groups, they do suggest a differential predominance of specific *pir* genes being upregulated in one disease group and less upregulated in the other disease group, as depicted in Table 2. At this stage, it remains challenging to assign precise functional implications to these fold-change differences; however, the observed trends indicate overall differences in *pir* gene expression profiles across the two disease groups.

The reason behind the differential expression of certain *pir* genes in the two disease groups could be associated with the host factors. Host factors significantly influence a parasite’s gene expression profile, a phenomenon driven by the dynamic interplay between the host’s internal environment and the parasite’s need to survive and reproduce. Key host factors include the host’s immune response, nutrient availability, age and sex, and the specific host tissue or organ the parasite infects (35–41).

### Principal component analysis

The PCA plot of all *pir* genes (Fig 5) from the microarray revealed a clear trend of partial segregation between the two severe malaria patient groups (CM, n=5; HDYS, n=12). While complete separation was not observed, distinct clustering patterns along PC1 and PC2 indicate that the cerebral malaria (n = 5) and hepatic dysfunction (n = 12) patient groups possess unique molecular signatures within the *pir* gene family. The variance explained by the PC1 and PC2 (28.6%) suggests that differential expression of *pir* genes contributes to the molecular heterogeneity associated with these severe malaria phenotypes.

**Fig 5.**
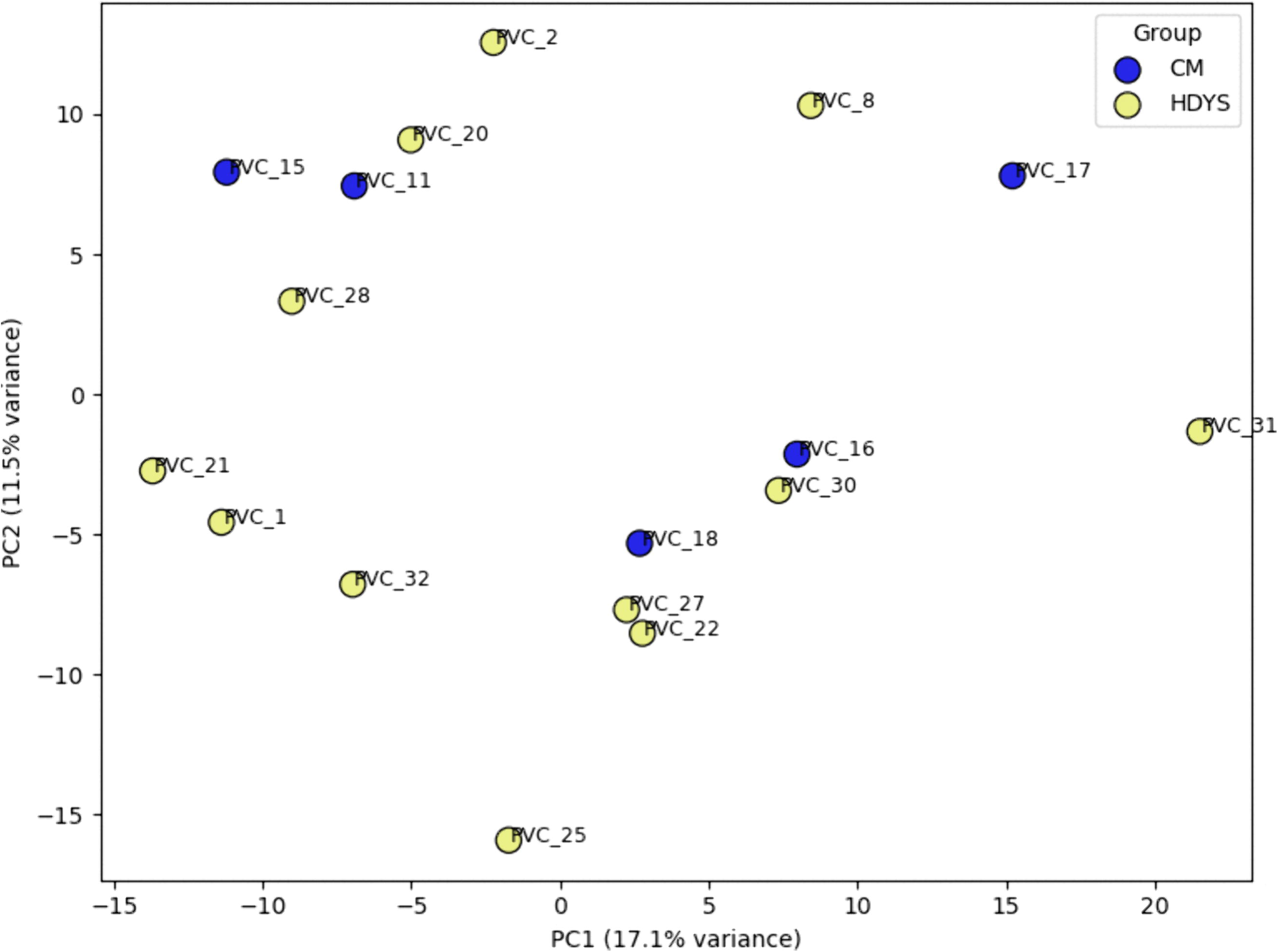
PCA plot of all the *pir* genes detected in the microarray. The comparison is made between 5 cerebral malaria patients against 12 hepatic dysfunction patients.

## Conclusion

In this study, we have reported the differences in the transcriptomic profile of *pir* genes in patients with *P. vivax* cerebral malaria and hepatic dysfunction. The upregulation of these *pir* genes may play a crucial role in pathogenesis. One of the possible roles of *pir* genes is that they may exhibit cytoadherence with host cell receptors, as reported in *Bernabeu et al* (23) and *Schappo et al* (24). An interesting observation of the study by *Bernabeu et al* (23) is that the *pir* gene documented to have shown binding, PVX_108770, *vir* C, in the reported study has shown at least 50% (HDYS n=6/12 to 12/12) upregulation in the patients’ group with hepatic dysfunction manifestation in our microarray data (Fig 2 and Table 2). Hepatic dysfunction in malaria is associated with heightened inflammation and apoptosis, particularly of Kupffer cells and lymphocytes. This provides a biologically plausible link between PIR-ICAM 1 interactions and HDYS pathogenesis (42) for which experimental approaches such as ELISA-based binding assays using recombinant PIR proteins and ICAM-1, streptavidin-biotin pull-down assays, and surface plasmon resonance (SPR) assays may be performed. Interestingly, *Fernandez-Becerra et al.* (35) have reported the cytoadhesion of the same *pir* gene product with ICAM-1 expressed by the human spleen fibroblasts cells in the spleen.

Different subcellular localisations of PIR proteins have also been shown, suggesting other possible functions of PIR proteins beyond cytoadherence (23). Functional assays, such as co-immunoprecipitation or interaction-based assays, can identify associated protein partners and provide insights into the additional roles of PIR proteins in parasite biology. These PIR proteins may also serve as important prognostic biomarkers for predicting disease progression. We acknowledge the limited sample size on which this analysis is based. However, samples showing these manifestations and correctly characterised are not easy to obtain. Further samples, when available, will be used to validate the data provided in this study. Additionally, the probable involvement of *pir* genes in causing severe disease requires further investigation.

## Supporting information

**S1 Table. Transcriptome profile of *pir* genes showing upregulation/downregulation in 5 CM and 12 HDYS patients’ group.**

**S2 Table. Comparative analysis of *pir* genes between CM and HDYS, upregulated/downregulated in at least 50% severe malaria patients’ group.**

**S3 Table. Distinct pir genes upregulated in one disease group (at least 50%) compared with their status in the other disease group.**

**S1 Fig. Details of primers used for quantitative real-time PCR analysis.** #Sequences are provided from 5’ to 3’ end. *All reactions were subjected to 35 cycles.

## Authors contribution

AD conceived the study, organised the material, and is the principal investigator of a project from the Department of Biotechnology and the Indian Council for Medical Research, India, and networked to facilitate the outcome. DKK and SKK performed the clinical investigations. AD, PAB, SS, and PR designed the experiments involved in this study. PAB performed RNA isolation, quality check and microarray data analysis. SS and PR performed the real-time qPCR experiment. SS and AD drafted the manuscript. SS, PR, and ST designed the tables, graphs and figures for this paper. PAB and SS are equal contributors to this work. All authors have read and approved the manuscript

## Competing interests

The authors have declared that no competing interests exist.

## Acknowledgements

We thank all the patients for their voluntary consent. We acknowledge BITS Pilani, Pilani campus and Sardar Patel Medical College for providing the required infrastructural facilities during this study. Ms Sheetal Middha and Jyoti Acharya were involved in the preprocessing of the samples at Sardar Patel Medical College, and their assistance is gratefully acknowledged. We thank the PlasmoDB (43) team for making the genome of the Sal I strain available. Genotypic Technology Pvt. Ltd., Bangalore, India, is acknowledged for the microarray hybridisation services provided.

## Financial Disclosure

AD is a Senior Professor at Birla Institute of Technology and Science (BITS) Pilani, Pilani campus, Rajasthan, India. SKK is a Senior Professor at Sardar Patel Medical College, Bikaner, Rajasthan. DKK is a former Senior Professor at Sardar Patel Medical College, Bikaner, Rajasthan. This study was partially funded by grants from Department of Biotechnology (DBT), New Delhi, India (https://dbtindia.gov.in/) through the grant BT/PR7520/BRB/10/481/2006 to AD, DKK and SKK and a grant from Indian Council for Medical Research (ICMR), New Delhi, India (https://www.icmr.gov.in/) through the Extramural Ad-hoc Scheme (Project ID: 2021-12909) to AD and SKK. The funders had no role in the study design, data collection and analysis, the decision to publish, or the preparation of the manuscript.

## Notes

### Competing Interest Statement

The authors have declared no competing interest.

### Summary of Updates

According to the reviwers comment, i have made the changes in the final manuscript.

## References

1. World malaria report 2025: Addressing the threat of antimalarial drug resistance.

2. Garrido-Cardenas JA, González-Cerón L, García-Maroto F, Cebrián-Carmona J, Manzano-Agugliaro F, Mesa-Valle CM. Analysis of Fifty Years of Severe Malaria Worldwide Research. Pathog Basel Switz. 2023 Feb 24;12(3):373.

3. Weiss DJ, Dzianach PA, Saddler A, Lubinda J, Browne A, McPhail M, et al. Mapping the global prevalence, incidence, and mortality of Plasmodium falciparum and Plasmodium vivax malaria, 2000–22: a spatial and temporal modelling study. The Lancet. 2025 Mar 22;405(10483):979–90.

4. Nauriyal D, Kumar D. Study of Severe Malaria Caused by Plasmodium Vivax in Comparison to Plasmodium Falciparum and Mixed Malarial Infections in Children. Biomed Pharmacol J. 2022 Sept 29;15(3):1597–604.

5. Phyo AP, Dahal P, Mayxay M, Ashley EA. Clinical impact of vivax malaria: A collection review. PLoS Med. 2022 Jan;19(1):e1003890.

6. Ozen M, Gungor S, Atambay M, Daldal N. Cerebral malaria owing to Plasmodium vivax: case report. Ann Trop Paediatr. 2006 June 1;26(2):141–4.

7. Thapa R, Patra V, Kundu R. Plasmodium vivax Cerebral Malaria. CASE Rep. 2007;44.

8. Genton B, D’Acremont V, Rare L, Baea K, Reeder JC, Alpers MP, et al. Plasmodium vivax and Mixed Infections Are Associated with Severe Malaria in Children: A Prospective Cohort Study from Papua New Guinea. PLOS Med. 2008 June 17;5(6):e127.

9. Kochar DK, Saxena V, Singh N, Kochar SK, Kumar SV, Das A. Plasmodium vivax Malaria - Volume 11, Number 1—January 2005 - Emerging Infectious Diseases journal - CDC. [cited 2024 Feb 3]; Available from: https://wwwnc.cdc.gov/eid/article/11/1/04-0519_article

10. Kochar DK, Pakalapati D, Kochar SK, Sirohi P, Khatri MP, Kochar A, et al. An unexpected cause of fever and seizures. The Lancet. 2007 Sept 8;370(9590):908.

11. Kochar DK, Tanwar GS, Khatri PC, Kochar SK, Sengar GS, Gupta A, et al. Clinical Features of Children Hospitalized with Malaria—A Study from Bikaner, Northwest India. Am J Trop Med Hyg. 2010 Nov 5;83(5):981–9.

12. Tanwar GS, Khatri PC, Sengar GS, Kochar A, Kochar SK, Middha S, et al. Clinical profiles of 13 children with Plasmodium vivax cerebral malaria. Ann Trop Paediatr. 2011 Nov 1;31(4):351–6.

13. Park SY, Park YS, Park Y, Kwak YG, Song JE, Lee KS, et al. Severe vivax malaria in the Republic of Korea during the period 2000 to 2016. Travel Med Infect Dis. 2019 July 1;30:108–13.

14. Guidelines for the treatment of malaria. Third edition | WHO | Regional Office for Africa [Internet]. 2025 [cited 2025 July 1]. Available from: https://www.afro.who.int/publications/guidelines-treatment-malaria-third-edition

15. del Portillo HA, Fernandez-Becerra C, Bowman S, Oliver K, Preuss M, Sanchez CP, et al. A superfamily of variant genes encoded in the subtelomeric region of Plasmodium vivax. Nature. 2001 Apr;410(6830):839–42.

16. Carvalho BO, Lopes SCP, Nogueira PA, Orlandi PP, Bargieri DY, Blanco YC, et al. On the Cytoadhesion of Plasmodium vivax–Infected Erythrocytes. J Infect Dis. 2010 Aug 15;202(4):638–47.

17. Janssen CS, Phillips RS, Turner CMR, Barrett MP. Plasmodium interspersed repeats: the major multigene superfamily of malaria parasites. Nucleic Acids Res. 2004 Oct 1;32(19):5712–20.

18. Carlton JM, Adams JH, Silva JC, Bidwell SL, Lorenzi H, Caler E, et al. Comparative genomics of the neglected human malaria parasite Plasmodium vivax. Nature. 2008 Oct;455(7214):757–63.

19. Lopez FJ, Bernabeu M, Fernandez-Becerra C, del Portillo HA. A new computational approach redefines the subtelomeric vir superfamily of Plasmodium vivax. BMC Genomics. 2013 Jan 16;14(1):8.

20. Auburn S, Böhme U, Steinbiss S, Trimarsanto H, Hostetler J, Sanders M, et al. A new Plasmodium vivax reference sequence with improved assembly of the subtelomeres reveals an abundance of pir genes. Wellcome Open Res. 2016 Nov 15;1:4.

21. Fernandez-Becerra C, Pein O, de Oliveira TR, Yamamoto MM, Cassola AC, Rocha C, et al. Variant proteins of Plasmodium vivax are not clonally expressed in natural infections. Mol Microbiol. 2005;58(3):648–58.

22. del Portillo HA, Lanzer M, Rodriguez-Malaga S, Zavala F, Fernandez-Becerra C. Variant genes and the spleen in *Plasmodium vivax* malaria. Int J Parasitol. 2004 Dec 1;34(13):1547–54.

23. Bernabeu M, Lopez FJ, Ferrer M, Martin-Jaular L, Razaname A, Corradin G, et al. Functional analysis of Plasmodium vivax VIR proteins reveals different subcellular localizations and cytoadherence to the ICAM-1 endothelial receptor. Cell Microbiol. 2012;14(3):386–400.

24. Schappo AP, Bittencourt NC, Bertolla LP, Forcellini S, da Silva ABIE, dos Santos HG, et al. Antigenicity and adhesiveness of a *Plasmodium vivax* VIR-E protein from Brazilian isolates. Mem Inst Oswaldo Cruz. 2022 Feb 4;116:e210227.

25. Bozdech Z, Mok S, Gupta AP. DNA Microarray-Based Genome-Wide Analyses of Plasmodium Parasites. In: Ménard R, editor. Malaria: Methods and Protocols [Internet]. Totowa, NJ: Humana Press; 2013 [cited 2024 Feb 3]. p. 189–211. (Methods in Molecular Biology). Available from: 10.1007/978-1-62703-026-7_13

26. Kafsack BFC, Painter HJ, Llinás M. New Agilent platform DNA microarrays for transcriptome analysis of Plasmodium falciparum and Plasmodium berghei for the malaria research community. Malar J. 2012 June 8;11(1):187.

27. Bubendorf L. High–Throughput Microarray Technologies: From Genomics to Clinics. Eur Urol. 2001 Oct 19;40(2):231–8.

28. Allison DB, Cui X, Page GP, Sabripour M. Microarray data analysis: from disarray to consolidation and consensus. Nat Rev Genet. 2006 Jan;7(1):55–65.

29. Boopathi PA, Subudhi AK, Middha S, Acharya J, Mugasimangalam RC, Kochar SK, et al. Design, construction and validation of a Plasmodium vivax microarray for the transcriptome profiling of clinical isolates. Acta Trop. 2016 Dec 1;164:438–47.

30. Das A, Holloway B, Collins WE, Shama VP, Ghosh SK, Sinha S, et al. Species-specific 18S rRNA gene amplification for the detection ofP. falciparumandP. vivaxmalaria parasites. Mol Cell Probes. 1995 June;9(3):161–5.

31. Pakalapati D, Garg S, Middha S, Kochar A, Subudhi AK, Arunachalam BP, et al. Comparative evaluation of microscopy, OptiMAL® and 18S rRNA gene based multiplex PCR for detection of *Plasmodium falciparum* & *Plasmodium vivax* from field isolates of Bikaner, India. Asian Pac J Trop Med. 2013 May 13;6(5):346–51.

32. Boopathi PA, Subudhi AK, Garg S, Middha S, Acharya J, Pakalapati D, et al. Revealing natural antisense transcripts from Plasmodium vivax isolates: Evidence of genome regulation in complicated malaria. Infect Genet Evol. 2013 Dec 1;20:428–43.

33. Untergasser A, Cutcutache I, Koressaar T, Ye J, Faircloth BC, Remm M, et al. Primer3—new capabilities and interfaces. Nucleic Acids Res. 2012 Aug 1;40(15):e115.

34. Fernandez-Becerra C, Yamamoto MM, Vêncio RZN, Lacerda M, Rosanas-Urgell A, Portillo HA del. Plasmodium vivax and the importance of the subtelomeric multigene vir superfamily. Trends Parasitol. 2009 Jan 1;25(1):44–51.

35. Fernandez-Becerra C, Bernabeu M, Castellanos A, Correa BR, Obadia T, Ramirez M, et al. Plasmodium vivax spleen-dependent genes encode antigens associated with cytoadhesion and clinical protection. Proc Natl Acad Sci. 2020 June 9;117(23):13056–65.

36. Host immune response against Plasmodium is shaped through host mediated alterations in parasite gene expression profile. | The Journal of Immunology | Oxford Academic [Internet]. [cited 2026 Jan 14]. Available from: https://academic.oup.com/jimmunol/article/212/1_Supplement/0486_5839/7995492

37. Cummings BE, Stucke EM, Coulibaly D, Lawton JG, Sobota RS, Kone AK, et al. Plasmodium falciparum gene expression signatures and antibody profiling implicate the fikk, phist, and surf multigene families in severe malaria syndromes. J Infect [Internet]. 2025 Dec 1 [cited 2026 Jan 9];91(6). Available from: https://www.journalofinfection.com/article/S0163-4453(25)00255-5/fulltext

38. Berland F, Bourret V, Peroz C, Malandrin L, Bonsergent C, Bailly X, et al. Effects of host sex, age and behaviour on co-infection patterns in a wild ungulate. Parasitology. 2025 Oct 1;1–15.

39. Ezenwa VO, Stefan Ekernas L, Creel S. Unravelling complex associations between testosterone and parasite infection in the wild. Funct Ecol. 2012;26(1):123–33.

40. Zuzarte-Luís V, Mota MM. Parasite Sensing of Host Nutrients and Environmental Cues. Cell Host Microbe. 2018 June 13;23(6):749–58.

41. Kumar M, Skillman K, Duraisingh MT. Linking nutrient sensing and gene expression in Plasmodium falciparum blood-stage parasites. Mol Microbiol. 2021;115(5):891–900.

42. Prenen F, Steen PEV den. Malaria-associated liver dysfunction: a forgotten challenge. Trends Parasitol. 2025 July 1;41(7):547–59.

43. Aurrecoechea C, Brestelli J, Brunk BP, Dommer J, Fischer S, Gajria B, et al. PlasmoDB: a functional genomic database for malaria parasites. Nucleic Acids Res. 2009 Jan 1;37(suppl_1):D539–43.

