## Supplementary figures and images for "Comparative clinical transcriptome of *pir* genes in severe *Plasmodium vivax* malaria"

### S1_Fig.tif

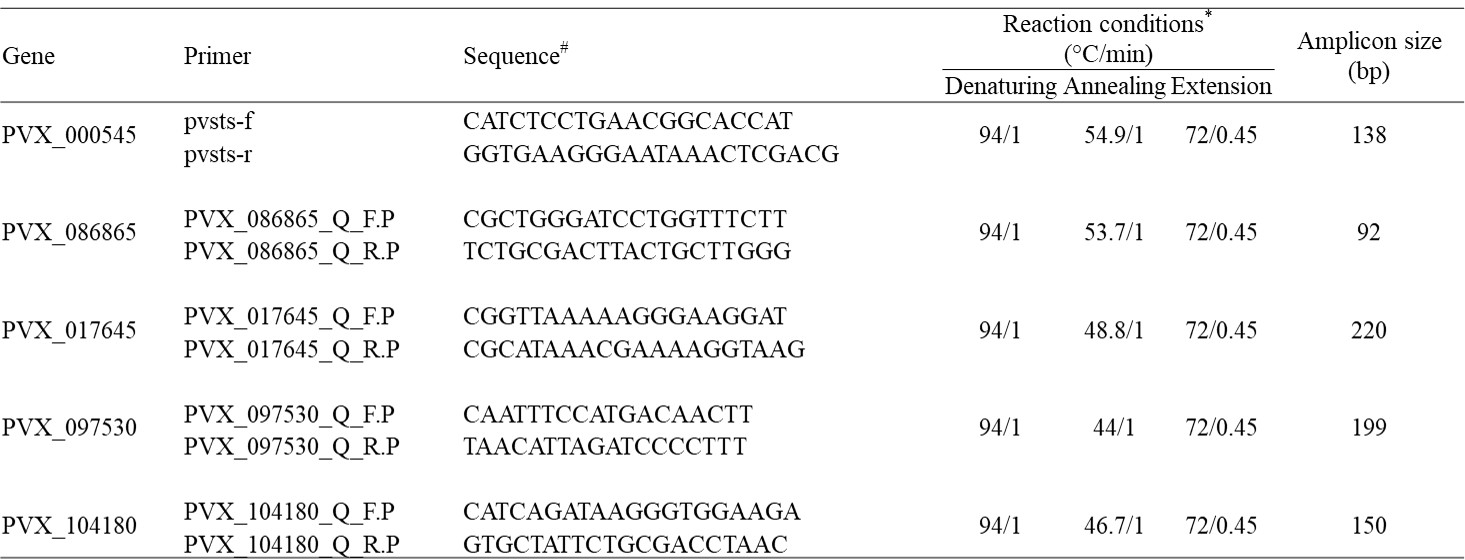
